# The *Plasmodium falciparum* artemisinin resistance-associated protein Kelch 13 is required for formation of normal cytostomes

**DOI:** 10.1101/2023.03.30.534478

**Authors:** Madel V. Tutor, Gerald J. Shami, Ghizal Siddiqui, Darren J. Creek, Leann Tilley, Stuart A. Ralph

## Abstract

Artemisinin (ART) is a quick-killing and effective antimalarial activated by the haem derived from haemoglobin digestion. Mutations in the parasite’s Kelch 13 (K13) protein compromise the efficacy of this drug. Recent studies indicate an undefined role for K13 in haemoglobin uptake. Here, we show that K13 is associated with the collar that constricts cytostomal invaginations required for the parasite to ingest host cytosol. Induced mislocalisation of K13 led to the formation of atypical invaginations lacking the cytostomal ring and constricted neck normally associated with cytostomes. Moreover, the levels of haemoglobin degradation products, haem and haemozoin, are decreased when K13 is inactivated. Our findings demonstrate that K13 is required for normal formation and/or stabilisation of the cytostome, and thereby the parasite’s uptake of haemoglobin. This is consistent with perturbation of K13 function leading to decreased activation of ART and consequently, reduced killing.

**Significance Statement:** Artemisinin-resistant parasites contain mutations in the gene encoding the Kelch 13 protein (K13). How K13 mutations result in artemisinin resistance is unclear. Here, we present evidence that normal K13 is required for the formation of the cytostome, a specialised parasite feeding apparatus used to endocytose host cell haemoglobin. Our results suggest that artemisinin resistance is due to a decrease in artemisinin activation brought about by a decrease in efficiency of haemoglobin uptake and consequently reduced production of haem.

## Introduction

Artemisinin and derivatives (ARTs) are sesquiterpene lactones with an endoperoxide bridge that can quickly and effectively kill the *Plasmodium* parasite during the pathogenic asexual blood stage. ARTs are administered in combination with other antimalarials (ART combination therapies, ACTs) due to artemisinin’s short *in vivo* half-life and to impede development of resistance (WHO, 2021). Unfortunately, resistance, characterised by a longer parasite clearance time, has emerged and spread throughout Southeast Asia, and more recently has arisen independently in Sub-Saharan Africa, where the majority of the disease burden exists (Balikagala et al., 2021; Uwimana et al., 2020). While mutations in some other genes are associated with ART resistance, mutations in the Kelch 13 (K13) gene (PF3D7_1343700 in the 3D7 strain) are strongly associated with resistance in the field and lead to the highest degree of resistance (Menard et al., 2016; Miotto et al., 2015; Straimer et al., 2015). Therefore, understanding the role of K13 can lead to a better understanding of how resistance arises and help inform changes in treatment regimen.

Recent studies indicate that K13 plays a role in haemoglobin uptake or endocytosis (Behrens et al., 2021 for review). While several studies have elucidated some of the key details of the haemoglobin uptake process, including its initiation at mid-ring stage, a putative role of actin in the transport process, and the identification of cytostomal invagination as the major route for uptake (Abu Bakar et al., 2010; Lazarus et al., 2008; Milani et al., 2015), considerable details remain to be described. Moreover, specific proteins important for this process and the cellular structures involved have not been unambiguously identified. Thus, it remains unclear how packets of haemoglobin cargo are transferred from the RBC cytosol to the digestive vacuole (DV) and how K13 is involved in this process (Spielmann et al., 2020 for review).

We have previously shown that K13 is associated with doughnut-shaped structures located at the parasite periphery (Yang et al., 2019). Recently, the *Toxoplasma* K13 orthologue was shown to have a similar localisation (Koreny et al., 2022). Using live-cell microscopy, we show that these *P. falciparum* K13 foci mark the positions at which the bulk of haemoglobin enters the parasite, supporting a role for K13 in haemoglobin uptake or endocytosis in the parasite. Moreover, we show that conditional disruption of K13 function disrupts formation of normal cytostomes and leads to a decrease in haemoglobin digestion products. We show that K13 is positioned at the electron dense collar around the neck of the cytostome and that its conditional mislocalisation disrupts the formation of this collar and the constriction of cytostomal invaginations; and blocks the generation of new cytostomes. Given its cellular location, we conclude that K13 is either a major constituent of, or is required for the generation of, the electron-dense cytostomal ring. K13 thus plays a crucial role in haemoglobin uptake through its function in the formation and stabilisation of cytostomal invaginations.

## Results

### K13 is associated with structures involved in uptake of host cytoplasm

Recent research on K13 points to a role in endocytosis (Birnbaum et al., 2020; Yang et al., 2019) and based on fluorescence imaging we previously proposed that GFP-tagged K13 localises to cytostomes (Yang et al., 2019). These double-membrane invaginations encompass both the parasitophorous vacuolar membrane (PVM) and the parasite plasma membrane (PPM), and serve as the “mouth” of the parasite for bulk uptake of the haemoglobin-containing host cytoplasmo (Abu Bakar et al., 2010; Milani et al., 2015). Electron microscopy reveals an electron dense cytostomal ring positioned immediately underneath the PPM and surrounding the narrowest part (neck) of the cytostomal invagination.

By deconvolution fluorescence microscopy (**Figure 1A)**, GFP-K13 is visualised as a punctate structure in ring stage parasites, with the number of puncta increasing in number as the parasite matures. As we have previously shown, these GFP-K13 puncta resolve as doughnut-shaped structures in super-resolution microscopy (Yang et al., 2019) (**Figure 1B**). Previous studies have shown that dextran, which is inert, stable, and commercially available conjugated to a range of fluorophores (Rivero & Maniak, 2006), can be taken up in cytostomal invaginations (Abu Bakar et al., 2010; Goodyer et al., 1997). To investigate the role of K13 in the uptake of red blood cell (RBC) cytosol, we resealed erythrocytes (as described by Abu Bakar et al., 2010 with modifications) in the presence of fluorescent dextran. We investigated the relationship of K13 puncta to cytostomes using time-lapse fluorescence microscopy.

**Figure 1.**
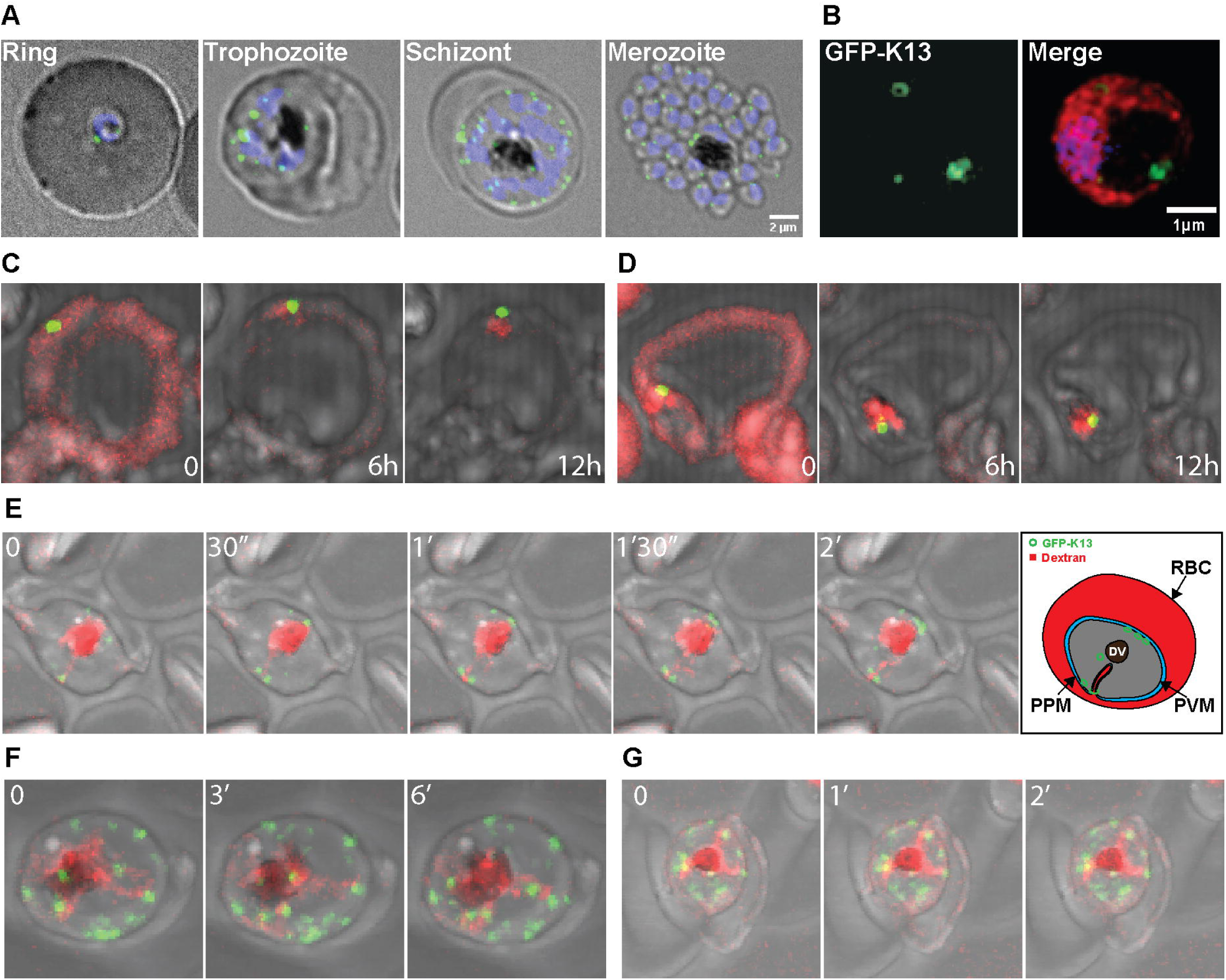
K13 is associated with sites of dextran uptake from the RBC. (A) Live-cell imaging of GFP-K13 transfectant-infected RBCs. K13 (GFP, green) and nucleus (DAPI-stained, blue) channels merged with DIC image. Fluorescence channels are displayed as maximum intensity projections while DIC is a single Z-slice. (B) Immunofluorescence assay (IFA) of GFP-K13-imaged using 3D structured illumination microscopy (3D-SIM). Parasites labelled with anti-GFP (K13-WT/, green), anti-GAPDH (cytoplasm, red), and DAPI (nucleus, blue). Maximum intensity projections are displayed; additional images in Figure S5. (C-H) Percoll-purified late-stage parasites cultured in resealed fluorescent dextran-loaded RBCs. (C, D) Fluorescence timelapse imaging of control early-stage parasite-infected RBCs. Timepoints illustrate the decrease of fluorescent dextran signal in the host compartment and its accumulation inside the parasite as the parasite matures. (E) Fluorescence time-lapse imaging of an untreated late-stage parasite-infected RBC. Yellow arrowheads indicate GFP-K13 associated with the peripheral end of a tubular fluorescent dextran structure. The last panel shows a cartoon representation of the 2’ panel. (F, G) Fluorescence timelapse imaging of JAS-treated late-stage parasite-infected RBCs. (F) Timepoints illustrate the repositioning of GFP-K13 together with the repositioning of the peripheral fluorescent dextran structure. (G) Late-stage GFP-K13-infected RBC showing three tubular fluorescent dextran structures, each associated at the peripheral end with a GFP-K13-labelled structure.

Transfected parasites expressing K13 N-terminally tagged with GFP (GFP-K13) (Birnbaum et al., 2017) were allowed to invade fluorescent dextran-loaded RBCs. Early-stage (∼10-22 hpi) or late-stage (∼30-42 hpi) parasites were prepared for 4D imaging as in Gruring & Spielmann, 2012 with modifications. As shown in **Figure 1 C, D, Movie S1, S2**, the fluorescent dextran was initially evenly distributed throughout the host compartment of the infected RBC, but in late ring stage becomes concentrated at a single point at the periphery of the parasite that coincides with the K13 punctum. These peripheral fluorescent dextran structures have previously been described (Abu Bakar et al., 2010) and concentrate the fluorescent cargo in a manner consistent with cytostomal invaginations. The co-localisation of the K13 punctum with the foci of dextran entering the parasite is consistent with K13 marking the position of the cytostome.

Of 50 cells scored, the GFP-K13 foci was found positively associated with a dextran structure for most of the duration of the experiment for 42 cells and for at least some of the duration in an additional 5 cells (**Figure S1**). Note that the microscopy mode used for these experiments is optimised for live cell imaging and is not compatible with the super-resolution optics required to resolve the K13 ring structure.

In late-stage parasites where the DV is fully formed (as evident from a visible mass of haemozoin crystals), dextran-containing structures are apparent as tubules originating from the periphery and extending or “snaking” towards the DV can be seen (**Figure 1E**). GFP-K13 (yellow arrowheads) can be seen associated with the peripheral extremity of these dextran structures. Additionally, as the tubule changes position over time, the GFP focus from which it originates moves so that it stays connected to the start of the tubule. This observation is consistent with the extended cytostomal invaginations in more mature parasites previously described by light and electron microscopy (Kennedy et al., 2019; Milani et al., 2015), and suggests that, as in the nascent cytostomes, GFP-K13 marks the point where the cytostomal invagination is constricted underneath the plasma membrane.

### An actin stabiliser does not inhibit K13-associated fluorescent dextran uptake

Actin plays a role in formation and trafficking of vesicles during endocytosis in other eukaryotes (Robertson et al., 2009); and actin-perturbation has been reported to disrupt cytostome formation in *P. falciparum* (Lazarus et al., 2008). We therefore investigated whether F-actin stabilisation might affect the K13-mediated uptake of host material. Parasites expressing GFP-K13 were cultured in fluorescent dextran-loaded RBC treated with 7 µM Jasplakinolide (JAS) for 3 hours, as per Lazarus et al., 2008, and imaged as described above. As shown in **Figure 1F-G**, JAS-treated parasites were still able to take up and concentrate the fluorescent-dextran in GFP-K13 associated structures. In late-stage JAS-treated parasites, we also see dextran tubules that are associated with the DV. Similarly, to untreated parasites, the peripheral terminus of these dextran structures is adjacent to a punctum of GFP-K13 (yellow arrowheads). This suggests that formation of the cytostomal ring is not readily disrupted by actin perturbation.

### Immunoelectron microscopy confirms K13 is localised to the cytostomal neck

As GFP-K13 is located at doughnut-shaped structures consistent with the dimensions of the cytostomal collar and is closely associated with haemoglobin uptake structures, we hypothesised that K13 plays a role in the function, maintenance and/or formation of the cytostome. To better define the localisation of K13 we performed immunoelectron microscopy using GFP-K13 parasites. Using anti-GFP antibodies, we found that the GFP-K13 fusion is found at the parasite-facing surface of the cytostomal neck, with labelling at or adjacent to the electron-dense cytostomal ring (**Figure 2A**). This localisation is consistent with the shape, dimensions, and position of K13 as determined using super-resolution fluorescence microscopy.

**Figure 2.**
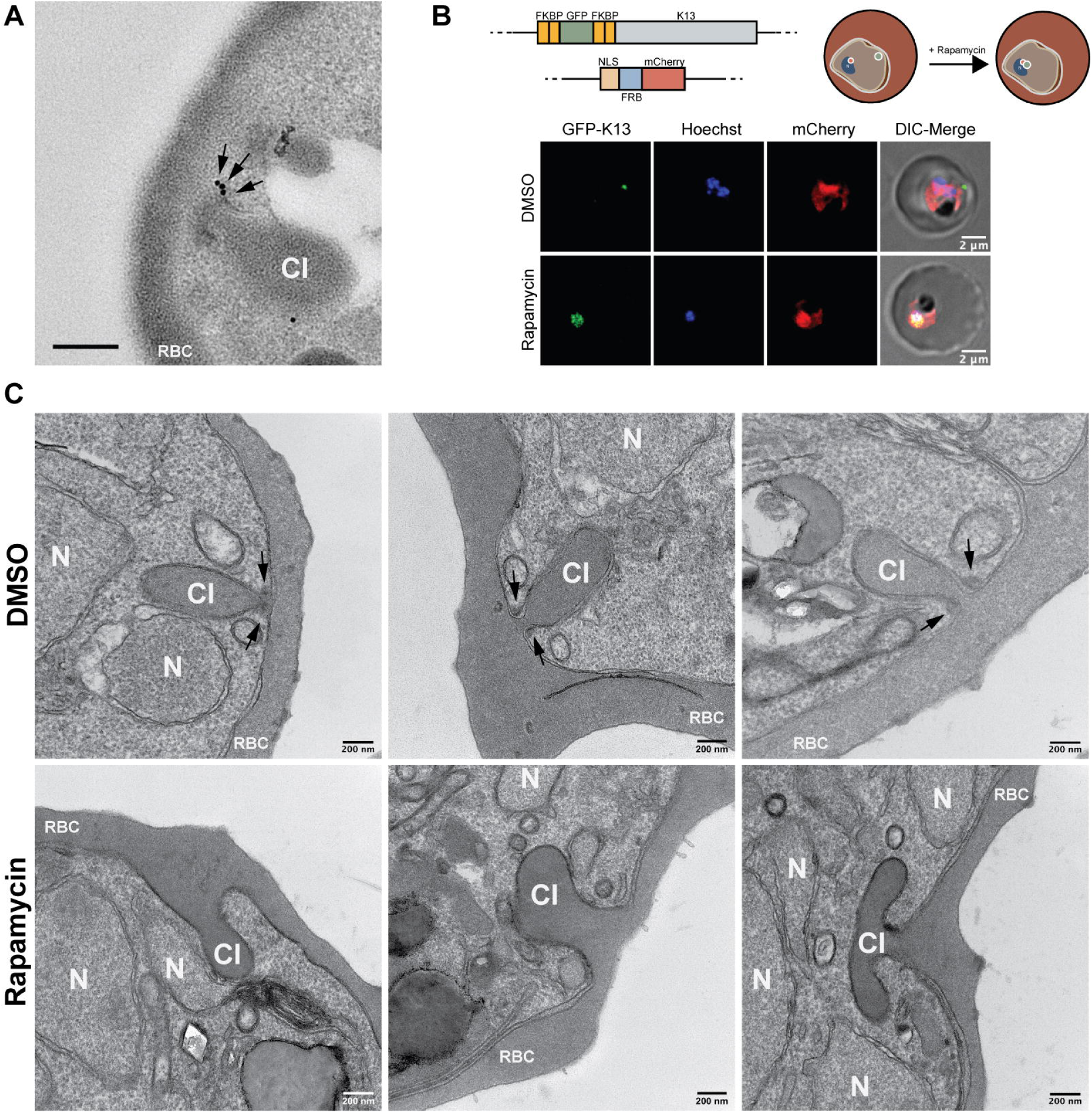
K13 is located at the cytostomal neck and its mislocalisation disrupts cytostome formation. (A) Immunoelectron micrograph of a late-stage GFP-K13 transfectant labelled with anti-GFP. Arrows indicate anti-GFP labelling on the neck of a cytostomal invagination (CI). Scale bar: 200 nm. (B, top panel) Molecular system for complete mislocalisation of GFP-K13. Schematic showing N-terminal-tagging of K13 with GFP and FKBP and the mislocaliser plasmid expressing NLS, FRB, and mCherry (Birnbaum et al., 2017). (B, top right) Cartoon representation of GFP-K13 mislocalisation. (B, bottom) Representative live-cell images showing mislocalisation of GFP-K13. Scale bar =2 µm. (C) Representative transmission electron microscopy (TEM) images showing invaginations in DMSO- and rapamycin-treated late-stage parasites. The top panel shows invaginations with shapes typical of a CI with electron-dense rings/collars around the tight necks. Arrows indicate electron dense cytostomal collars. Bottom panel shows invaginations with irregular shapes and wide necks lacking the electron-dense rings/collars. N: nucleus.

### K13 mislocalisation disrupts cytostome formation

To characterise the impact of K13 loss of function on the cytostome, we used a knock-sideways (KS) parasite line that allows complete conditional inactivation of K13 (K13-100KS) by protein mislocalisation (Birnbaum et al., 2017) (**Figure 2B**). Here, we examined the effects of the complete inactivation of K13 in the first cycle. We first aimed to identify an appropriate timing for initiating the mislocalisation to obtain completely mislocalised but not growth-arrested parasites. Complete mislocalisation of K13 allowed completion of that cycle and led to ring-stage arrest in the next cycle (**Figure S2-S3, Table S1**), consistent with a previous report (Birnbaum et al., 2017). We therefore initiated mislocalisation treatment in early-stage parasites (∼10-22 hpi) and harvested 24 hours later (∼30-42 hpi). Following a 24-hour treatment initiated at early-stage, the harvested late-stage parasites were prepared for imaging by electron microscopy (EM). We focused our attention on trophozoite-stage parasites because cytostomes are easiest to find during this stage, and because parasites could be magnet-purified for preparation of EM samples. Transmission electron microscopy (TEM) of infected RBCs in 250 nm sections showed that in late-stage parasites where K13 was mislocalised, maintenance or formation of cytostomes was perturbed (**Figure 2C**). While the cytostomes in control cells exhibited a characteristic constricted neck, surrounded by an electron-dense collar, both the constriction and the electron dense collar were absent in K13-mislocalised parasites. While untreated parasites exhibited elongated invagination that extended into the cell, K13-mislocalised parasites exhibited shallow invaginations that were flattened at the parasite periphery or branched immediately inside the parasite.

### Tomographic characterisation of invaginations in K13 mislocalised parasites

To allow observation of a greater number of cells and 3-dimensional analyses of the cytostomal invaginations, we performed serial block face scanning electron microscopy (SBF-SEM) on untreated and K13-mislocalised parasites using 50 nm slices. This allowed the complete 3D inspection of hundreds of cells in each condition (**Figure 3A-B, Movie S3, S4**) and revealed that cytostomal number and structure were dramatically different between untreated and K13-mislocalised parasites (**Figure 3C**). Three-dimensional reconstruction of three of these cells for each of the DMSO-(carrier only) and rapamycin-treated (mislocalised) samples was performed, and the rendered models are presented in **Figure 4A**. At the time of commencement of K13 mislocalisation, *i*.*e*., ring stage, parasites possess a single cytostome (Hanssen et al., 2013). Over the 24-hour treatment, *i*.*e*., during trophozoite maturation, the untreated parasites form additional cytostomes (Abu Bakar et al., 2010; Milani et al., 2015). However, in parasites with mislocalisation of K13, the cytostomes that were evident had an altered shape, with a loss of the constriction at the cytostome neck (**Figure 4A**) and most cells completely lacked a normal cytostomal invagination. This suggests that formation of new cytostomes and maintenance of existing cytostomes was perturbed. We selected 100 mid- to late-trophozoite stage parasites that were single infections that had not undergone segmentation (*i*.*e*., schizont maturation), for both untreated and rapamycin-treated K13-100KS parasites. The cytostomes, defined as invaginations with a narrow neck and open to the host cytoplasm, were counted in each of the selected parasites. 78 untreated parasites contained a cytostome while only 32 rapamycin-treated parasites contained a cytostome. 11 untreated parasites compared to only three rapamycin-treated parasites contained more than one cytostome. In 200 parasites scored, we saw a 59% decrease in the number of trophozoite-stage parasites with normal cytostomes. We noticed no other visible aspects of cellular morphology, organelle structure, or apparent parasite maturity that differed between the untreated and K13-mislocalised parasites.

**Figure 3.**
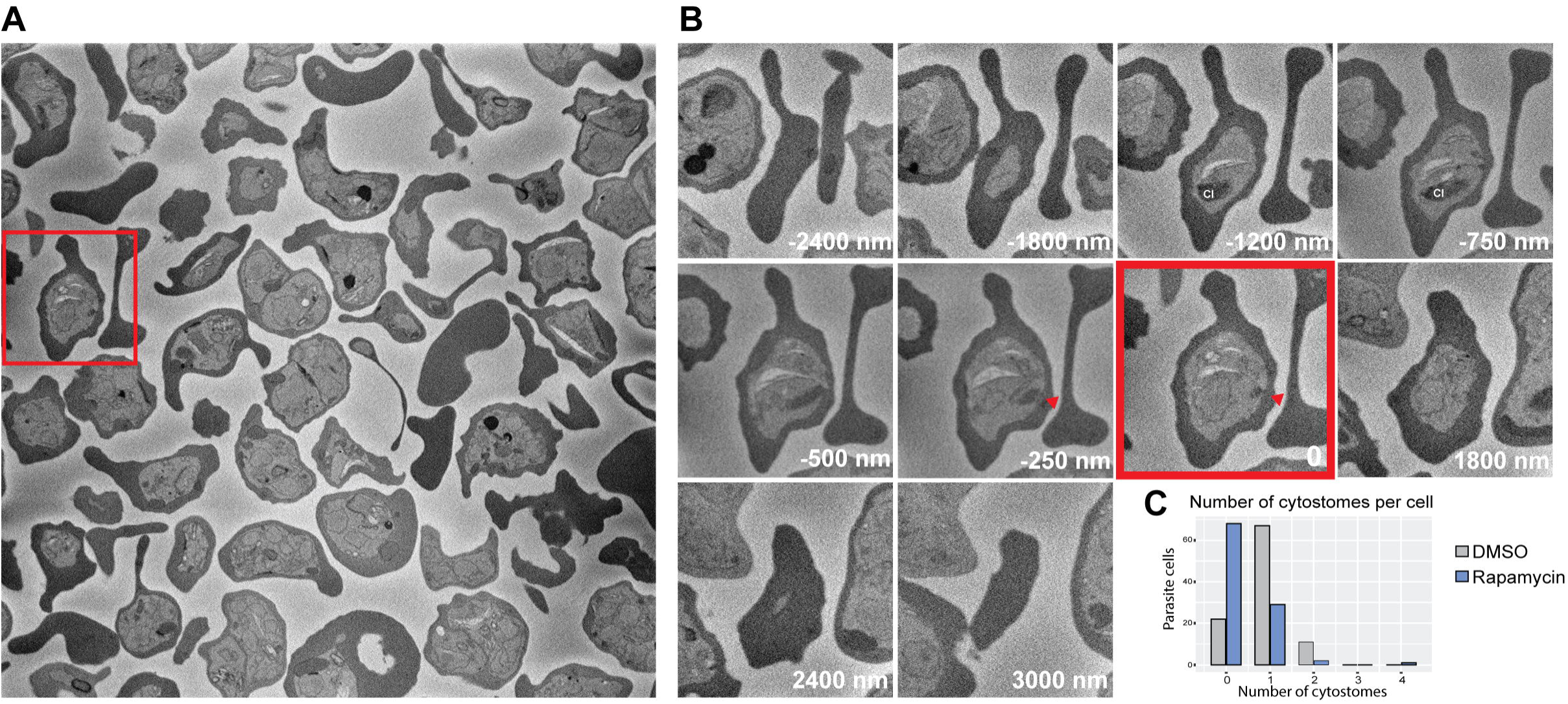
Quantitative analysis of atypical invaginations formed upon mislocalisation of K13. Representative single plane Serial Block Face Scanning Electron Microscopy (SBF-SEM) image (section thickness = 50 nm) showing many DMSO-treated K13-100KS-infected RBCs. Additional slices along the Z-axis of the highlighted infected RBC in A (red box). Red arrowhead points to opening of the CI to the RBC cytosol. (C) Comparison of the number of normal cytostomes formed in DMSO- and rapamycin-treated K13-100KS-infected RBCs. Normal cytostome is defined as an invagination with a tight neck in mid to late trophozoites that have not segmented and present as single infections. x-axis: number of cytostomes; y-axis: number of infected RBCs. 100 infected RBCs were counted per treatment.

**Figure 4.**
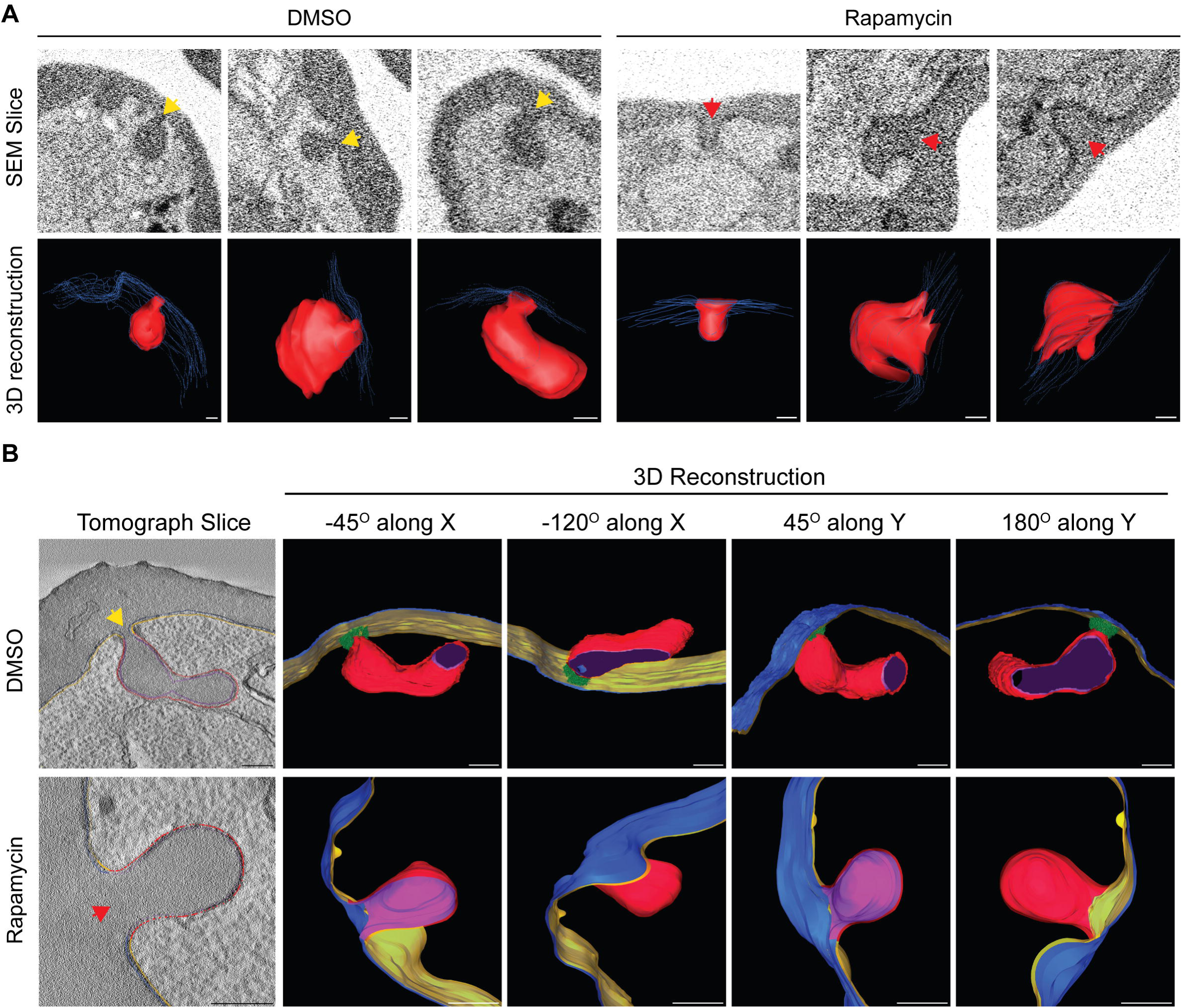
Tomographic analysis of cytostomal invaginations upon mislocalisation of K13. (A) 3D reconstructions of serial block-face scanning electron microscopy (SBF-SEM) images of DMSO- and rapamycin-treated late-stage K13-100KS-infected RBCs. Yellow arrowheads point at cytostomal invaginations with a tight neck opening to the RBC cytosol. Red arrowheads point at cytostomal invaginations with a wide neck opening to the RBC cytosol. (B) 3D reconstructions of transmission electron tomography images (section thickness = 250 nm; tilt range = ± 60°) of DMSO- and rapamycin-treated late-stage K13-100KS-infected RBCs. Yellow: parasitophorous vacuolar membrane (PVM); Blue: parasite plasma membrane (PPM); Red: PVM surrounding cytostomal invagination; Dark purple: PPM lining cytostomal invagination in DMSO-treated parasites; Light purple: PPM lining cytostomal invagination in rapamycin-treated parasites; Green: electron-dense collar around cytostomal neck; Yellow arrow: invagination with a tight neck surrounded by an electron-dense collar; red arrow: invagination with a wide neck lacking an electron-dense collar. Scale bars = 200 nm.

To further characterise the cytostome phenotype, we conducted transmission electron tomography on semi-thin slices (250 nm thick) imaged through a rotational tilt range of ±60°. Reconstructed cytostomes from untreated or K13-mislocalised parasites are shown in **Figure 4B and Movie S5, S6**. This analysis confirmed that in the untreated samples, parasites cytostomes were constricted into a tight neck at the point where the invagination meets the PV (with an internal diameter of the PM invagination of 55 nm and surrounded by an electron-dense collar) (**Supp fig S4**). However, in the K13-mislocalised parasites, although invaginations are present, the constriction around the invaginations is limited or absent, and no electron dense collar is present. Together the 3D electron microscopy data indicate that K13 is required for the formation or maintenance of the electron dense cytostomal collar, and that in the absence of this collar, cytostomal morphology and biogenesis or stability is disrupted.

### K13 mislocalisation leads to a decrease in haemoglobin digestion products

Most haemoglobin digestion occurs inside the DV. During this process, haemoglobin is degraded into large fragments, liberating the prosthetic haem group, which is toxic in its free form. To circumvent toxicity from haem, the parasite biocrystalises haem into haemozoin. Previous studies have shown that haemoglobin uptake and catabolism is reduced when K13 is mutated or is partially mislocalised (Birnbaum et al., 2020; Siddiqui et al., 2017; Yang et al., 2019) and these data are consistent with our observation that complete K13 mislocalisation perturbs the cytostome ultrastructure. We hypothesised that complete K13 mislocalisation would also lead to reduced production of the haem and haemozoin by-products of haemoglobin digestion. Decreased production of free haem is proposed to underpin ART resistance; however, to our knowledge, it has not been measured in K13 knock-sideways parasites.

Mislocalisation of K13 was induced and parasites were harvested as described above. Following a protocol adapted from Combrinck et al. and Birrell et al. fractionation of haem-containing molecules was performed (**Figure 5A**), and levels of free haem and hemozoin were measured using a colorimetric assay (Birrell et al., 2020; Combrinck et al., 2015). Mislocalisation of K13 caused a significant decrease in the level of free haem in both early- (p = 0.0166) and late-stage (p = 0.0005) parasites, as well as a significant decrease in haemozoin (p = 0.0093) in late-stage parasites (**Figure 5B**). This provides direct evidence that disruption of normal K13 function leads to a reduction in the abundance of haem that is required for ART activation.

**Figure 5.**
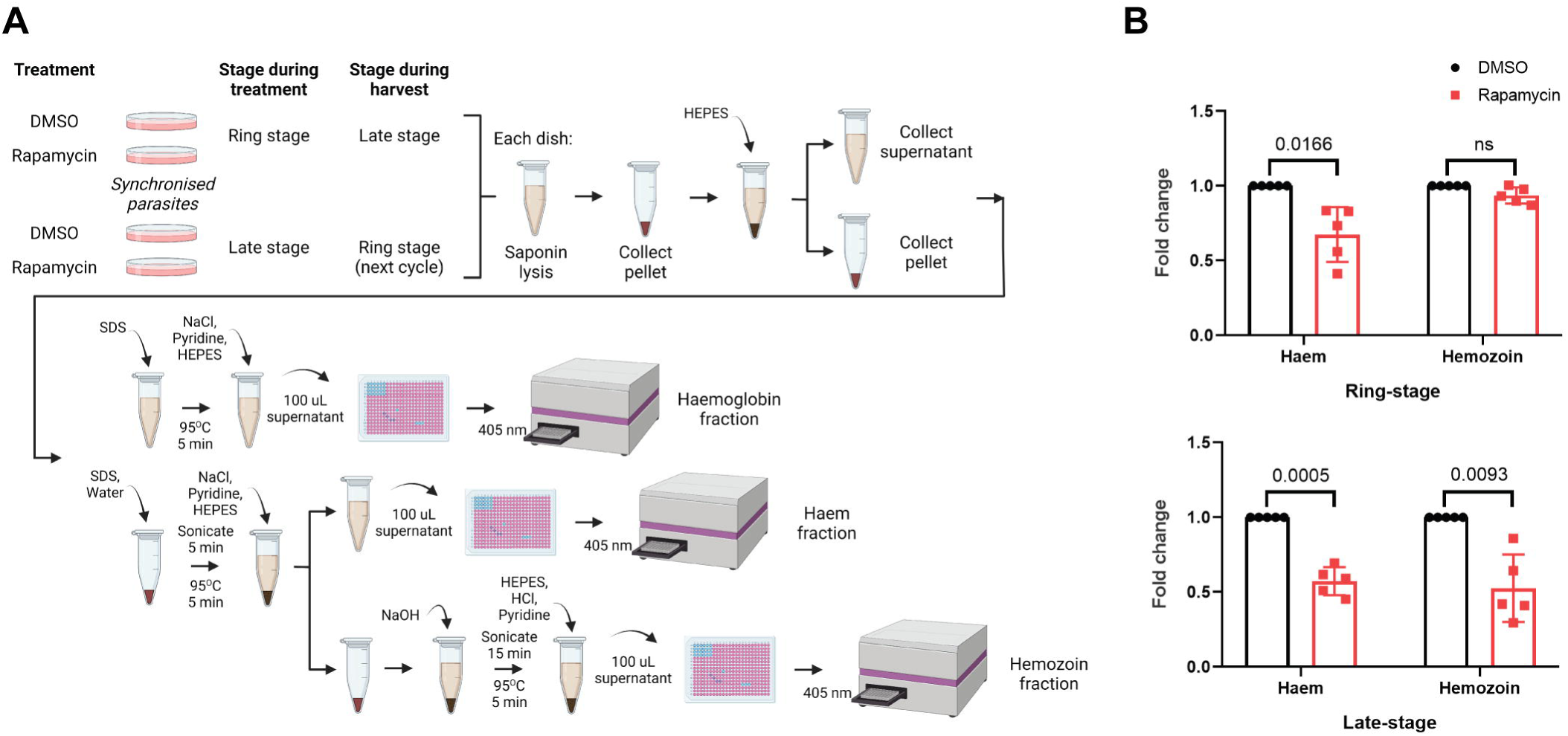
K13 mislocalisation decreases free haem in ring- and late-stage, and haemozoin in late-stage, parasites. (A) Mislocalisation and haem fractionation workflow. Created with BioRender.com. (B) Graphs showing effect of K13 mislocalisation on haem and haemozoin levels. Top graph shows results from parasites harvested at early stage (next cycle), showing a significant decrease in the free haem fraction (p = 0.0166) in K13-mislocalised parasites compared to untreated controls. Bottom graph shows results from parasites harvested at late stage, showing a significant decrease in the haem (p = 0.0005) and haemozoin (p = 0.0093) fractions in K13-mislocalised parasites (n = 5).

## Discussion

Recent published studies indicate that K13 plays a role in haemoglobin uptake or endocytosis but the mechanism by which K13 participates is unclear (Birnbaum et al., 2020; Koreny et al., 2022; Siddiqui et al., 2017; Yang et al., 2019). The doughnut-shaped structures we observed in super-resolution images of GFP-K13 transfectants suggested a role in the function of the cytostomal collar. Here we investigate that possibility using light and electron microscopy. Using live-cell imaging of parasite uptake of host cytosol, we find that GFP-K13 is located at the site of host cytosol uptake, consistent with GFP-K13 residing at the parasite cytostomal collar. This is confirmed by our localization using immuno-electron microscopy of GFP-K13 to the electron dense cytostomal neck. Our multiple ultrastructural investigations also demonstrated that conditional inactivation of GFP-K13 led to failure to form normal cytostomal invaginations, expansion of the cytostomal neck, and the disappearance of the electron-dense cytostomal collar. Disrupted cytostome formation, in turn, led to a significant decrease in haemoglobin degradation products (haem and haemozoin) in both the ring- and late-stage parasites. Together, these data suggest that K13 is a component of the electron dense cytostomal collar and is essential for the formation or maintenance of this structure. The recent report that K13 forms a higher order oligomeric structure is also consistent with the possibility that K13 is a major repeating structural element in the cytostomal collar (Goel et al., 2022).

### Cytostomal collar

Several studies have reported that the active cytostome is the major route of haemoglobin uptake in the parasite. Although a cytostomal invagination is not apparent in merozoites, a cytostomal ring is present (Bannister et al., 2004). Thus, each newly invaded ring stage parasite has a cytostomal ring ready to initiate formation of cytostomal invaginations (Hanssen et al., 2013; Milani et al., 2015). Inhibition of the uptake pathway in *P. falciparum* has been shown to lead to formation of long haemoglobin-filled tubes devoid of the electron-dense collar at the neck and to accumulation undigested haemoglobin ultimately resulting in parasite death (Milani et al., 2015). This is consistent with the cytostomal collar playing an essential role in the parasite via its function in the formation or maintenance of the cytostome.

The process that initiates formation of the cytostomal invagination at the K13 ring is unclear. It may be driven by a direct interaction of the cytostomal collar with the parasite plasma membrane or may require additional force exerted by polymerisation of actin potentially driven by myosin C. In clathrin-mediated endocytosis in mammalian cells, clathrin coats can spontaneously assemble but interaction with the cargo and other associated proteins are needed to stabilise the structure (Kumari et al., 2010). However, cytostomes do not microscopically resemble clathrin-coated pits and endocytosis appears to function independently of clathrin (Birnbaum et al., 2020; Henrici et al., 2020). Interestingly, however, other typical clathrin-associated proteins have been shown to associate with K13 namely Eps15, UBP1, and AP-2µ (Birnbaum et al., 2020; Henrici et al., 2020; Henriques et al., 2013; Henriques et al., 2015), and mutations in these genes are also associated with artemisinin resistance. Likewise in *Toxoplasma*, the K13 homologue interacts with many parts of the endocytic machinery, but not with clathrin (Koreny et al., 2022). Further work is warranted to dissect the mechanism of such a divergent, clathrin-independent endocytocytic pathway.

Our live-cell imaging of parasite feeding indicates a transition from early stages, where K13 marks the site of a small focus of host cytosol, to later stages that contain a distinct tubule initiating at the K13-defined cytostome. We observed these structures in untreated and Jasplakinolide-treated cells. These tubular structures have previously been described in mature parasites (Elliott et al., 2008; Kennedy et al., 2019; Milani et al., 2015) – although in one study only in the presence of Jasplakinolide. The discovery that these tube structures initiate at the site of a K13 focus, and that the K13 focus repositions together with the cytostomal structures reinforces this as an important aspect of haemoglobin uptake. By electron microscopy we did not see these tubular structures connect directly to the DV (though they are closely located), and the means by which trafficking between the tubule and the DV occurs requires further study. Milani and colleagues have previously investigated the function of the electron-dense collar in the fission of the invagination from the membranes. They found no evidence that the collar has a role in this process and proposed that it is likely involved in the stability of the cytostome (Milani et al., 2015). Our finding that cytostomal rings are always approximately the same size, across many parasites, is consistent with these being stable structures.

Our imaging reveals that the formation of normal cytostomes is disrupted in parasites when K13 was conditionally inactivated. K13 must therefore play a role in this process, possibly by initiating formation of the collar, as its localisation in the potential sites of invagination appear to precede the appearance of the invagination. That is, live-cell imaging revealed some GFP-K13 puncta alone at the periphery prior to the appearance of a visible focus of ingested dextran. Based on the absence of cytostomes with a tight neck in the K13-mislocalised parasites, it is also possible that K13 plays a role in the maintenance of the integrity of the invagination, perhaps preventing the structure from collapsing.

Lastly, we show that K13 mislocalisation leads to a decrease in haemoglobin degradation by-products that are relevant to artemisinin activation, further supporting the role of K13 in artemisinin activation and resistance. Previous studies reported a decrease in peptides that are intermediates of haemoglobin degradation when K13 is 90% mislocalised and in K13 mutants consistent with our findings (Siddiqui et al., 2017; Yang et al., 2019).

### K13 and ART resistance

If K13 is required for normal cytostome formation, how do K13 mutations mediate artemisinin resistance? One possibility is that mutations compromise K13’s ability to form a cytostomal ring. However, the most prevalent K13 mutations have no apparent effect on the ability of K13 to oligomerise (Goel et al., 2022). Other studies point to K13 mutations primarily impacting the parasite through modifying the overall abundance of K13 protein (Birnbaum et al., 2020; Siddiqui et al., 2017; Yang et al., 2019). K13 mutations lead to lower abundance of K13 and thus may result in fewer cytostomal rings and therefore fewer cytostomes. It is also possible that the K13 mutants are still able to form the same number of cytostomes, but the mutations lead to unstable, smaller, or faulty cytostomes, with reduced endocytic function. In our experiments, the complete mislocalisation of K13 from the cytostome led to the formation of unusual invaginations with wider necks and atypical shape, as well as a decrease in haem that activates ART. These parasites are unlikely to simulate the exact phenotype of an ART resistant mutant with decreased K13, but nonetheless suggests resistance mutations may impact cytostome function. It remains a mystery why there are numerous mutations that lead to a decrease in K13 abundance and confer ART resistance, but no parallel reports of mutations causing changes in K13 transcript abundance.

Overall, these results provide an explanation as to how K13 could be involved in haemoglobin uptake or endocytosis in the parasite. In the context of ART resistance, disruption in the formation of the cytostome leads to a decrease in haemoglobin taken up by the parasite. This results in less haem being available to activate ART, thereby allowing the parasite to survive ART pressure. Mutations in K13 that produce viable parasites must still allow the parasite to establish sufficient uptake structures that they can adequately support the growth of the parasite. Given the lethality of total functional disruption of K13 (K13-100KS), and K13’s essentiality in other studies, mutations in K13 that lead to complete loss of cytostomal function must be lethal. The precise molecular function of K13 and how K13 mutations impact cytostome function and haemoglobin uptake promise to be fruitful topics of further investigation.

## Materials and methods

### *P. falciparum* in vitro culture

*P. falciparum* 3D7 GFP-K13 and K13-100KS (Birnbaum et al., 2020) parasites were cultured as described in Trager & Jensen, 1976 and were sorbitol or Percoll-synchronised as described in (Radfar et al., 2009). Magnet purification was done following the method of (Kim et al., 2010).

### K13 mislocalisation

Sorbitol-synchronised ∼5% ring-stage parasitemia 2% haematocrit K13-100KS cultures (Birnbaum et al., 2017) were Percoll-purified after ∼24 hrs, harvesting late-stage parasites. Harvested pellets were then split into two cultures and maintained with 0.9 mM DSM-1. After ∼24 hours, one culture was treated with 250 nM rapamycin (Cayman Chemical, 13346) and another with 100% DMSO (equal volume to rapamycin). Treatment for early-stage (∼10-22 hpi) harvest is initiated during late-stage (∼30-42 hpi) while for late-stage harvest, treatment was initiated during early-stage. Parasites for EM or haem fractionation were harvested ∼24 hours post-treatment initiation.

### Live-cell and immunofluorescence assay microscopy

Single timepoint live-cell microscopy was performed by applying 5 µL, 4% haematocrit Hoechst 33342-stained DMSO- or rapamycin-treated K13-100KS cultures onto a slide, covered with a coverslip, and sealed with nail polish. Imaging was performed using DeltaVision DV Elite deconvolution microscope (Applied Precision) using a 100X U-Plan S-Apo oil immersion objective. The images were deconvolved using SoftWoRx 6.1 and analysed in Fiji (Schindelin et al., 2012).

For timelapse live-cell microscopy, dextran-loaded RBCs were prepared using either SNARF-1 10,000 MW (Invitrogen™, D3303) or tetramethylrhodamine 10,000 MW, lysine-fixable (Invitrogen™, D1817) following a protocol adapted from Abu Bakar et al., 2010. Purified late-stage asexual GFP-K13 parasites were grown in dextran RBCs. Following adapted protocol from (Gruring & Spielmann, 2012), parasites were immobilised using PHA-E (Sigma, L8629) on a 35 mm high µ-Dish (ibidi, 81156). Sterile PBS-washed pellet of GFP-K13 parasites resuspended in PBS (<1% haematocrit) was allowed to settle onto washed µ-Dish at 37^°^C. Unattached cells were washed off with PBS and the dish was filled with prewarmed complete media (∼8 mL). The dish was locked and sealed with parafilm. Images were taken using Zeiss LSM880 Elyra with the Alpha Plan-Apochromat 100X oil immersion objective using the confocal detectors (GaAsP and PMT). The system was prewarmed to 37^°^C for at least 2 hours prior to usage. Images were analysed and processed in Fiji (Schindelin et al., 2012).

Immunofluorescence assay was performed on PHA-E-immobilised parasites by fixation with 2% paraformaldehyde and 0.008% glutaraldehyde (Electron Microscopy Sciences) and permeabilisation with 0.1% Triton X-100 in PBS. Primary antibodies used were mouse anti-GFP (1:300, Roche 11814460001 Anti-GFP, from mouse IgG1K (clones 7.1 and 13.1)) and rabbit anti-GAPDH (1:300, (Jackson et al., 2007)). Secondary antibodies used were anti-mouse or anti-rabbit conjugated with AlexaFluor® 488 or 568, respectively. Antibodies were prepared in 3% BSA in PBS. 3D structure illumination microscopy (3D-SIM) was done using DeltaVision OMX V4 Blaze (GE Healthcare) using 60X Olympus Plan APO N oil immersion objective. Images were processed using Fiji (Schindelin et al., 2012).

### Electron microscopy

Magnet-purified parasites were fixed in 2% glutaraldehyde and 2% paraformaldehyde in 0.1 M Sorensen’s Phosphate Buffer (SPB) and post-fixed in 1% osmium tetroxide and 1.5% potassium in 0.1 M SPB. Samples were incubated in 1% tannic acid and 1% uranyl acetate to enhance cellular contrast. Samples were dehydrated in an ascending solvent series, progressively infiltrated with hard-grade Procure 812 resin and polymerised at 60°C for 48 h.

For TEM and TET, sections were generated using a Leica EM UC7 ultramicrotome (Leica Microsystems) and collected on 200-mesh Cu and single slot grids, respectively. Sections were post-stained using aqueous uranyl acetate and lead citrate for 10 min each. Ultrathin sections were imaged on a FEI Tecnai G2 F30 at 200 kV. TET was performed on semithin sections over a tilt range of ± 60°. Tomograms were aligned and reconstructed using Etomo and segmented and visualised using 3dmod (bundled with the IMOD software package) (Kremer et al., 1996).

For SBF-SEM, samples were mounted onto an aluminium pin, dried, and coated with 10 nm layer of gold. Sequential inverted backscattered electron images were acquired every 50 nm using a Teneo VolumeScope (FEI Company) at 3 kV under low vacuum conditions. Images were processed using Fiji. Image binning, histogram normalisation, and alignment using the StackReg plugin were done in Fiji (Schindelin et al., 2012). Regions of interest were segmented and visualised using 3dmod (Kremer et al., 1996).

For immunoelectron microscopy, samples were fixed in 1% glutaraldehyde, dehydrated in an ethanol series, and embedded in LR white resin. Ultrathin sections (∼100 nm) were labelled using mouse anti-GFP antiserum (1:300, Roche 11814460001) and 18 nm Colloidal Gold AffiniPure Goat Anti-Mouse (Jackson, 115-215-166) and post-stained with 2% uranyl acetate and lead citrate. Imaging was performed on a FEI Tecnai F20.

### Haemoglobin, haem and haemozoin fractionation assay

*P. falciparum* 3D7 GFP-K13 and K13-100KS (Birnbaum et al., 2017) parasites induced for mislocalisation as described above were saponin-lysed to harvest pellets for measuring haemoglobin, haem, and haemozoin levels using an adapted protocol from Birrell et al., 2020 and Combrinck et al., 2015. Data analysis was conducted using GraphPad Prism 8.0 and MS Excel. Raw and processed data are available in **Table S2-S3**. Significance was calculated using one sample t-test (hypothetical value: 1) on fold-change values relative to untreated (DMSO) control.

## Acknowledgements

We gratefully thank the Ian Holmes Imaging Centre and the Biological Optical Microscopy Platform for their support and assistance in this work. We thank the Australian Red Cross service for blood donations. We thank Dr. Tobias Spielmann and Dr. Jakob Birnbaum, Bernhard Nocht Institute for Tropical Medicine, for GFP-K13 and K13-100KS parasites.

