## Supplemental Figures 1 for "The *Plasmodium falciparum* artemisinin resistance-associated protein Kelch 13 is required for formation of normal cytostomes"

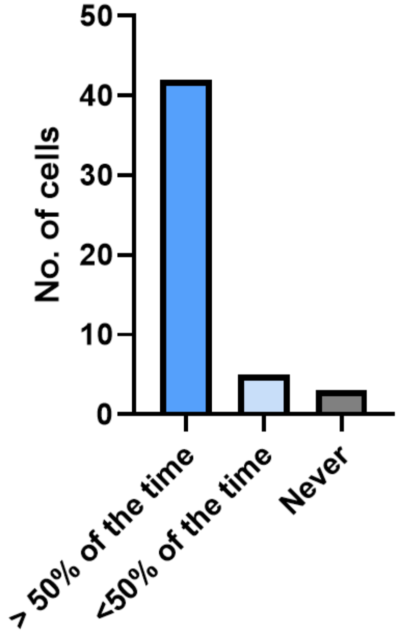


**Figure S1.** Quantification of GFP-K13/fluorescent dextran live-cell timelapse imaging. Fifty untreated cells were selected based on whether the imaging was stable with respect to all axes, the angle/view of the parasite was full, and the shape/morphology was distinguishable for all timepoints. Only parasites from early ring to late trophozoite stages were considered. Parasites were scored as "> 50% of the time" for cells with at least one GFP-K13 near dextran structures >50% of the time, "<50% of the time" for cells with at least one GFP-K13 near dextran structures <50% of the time, and "Never" for cells where GFP-K13 was never seen near a dextran structure.

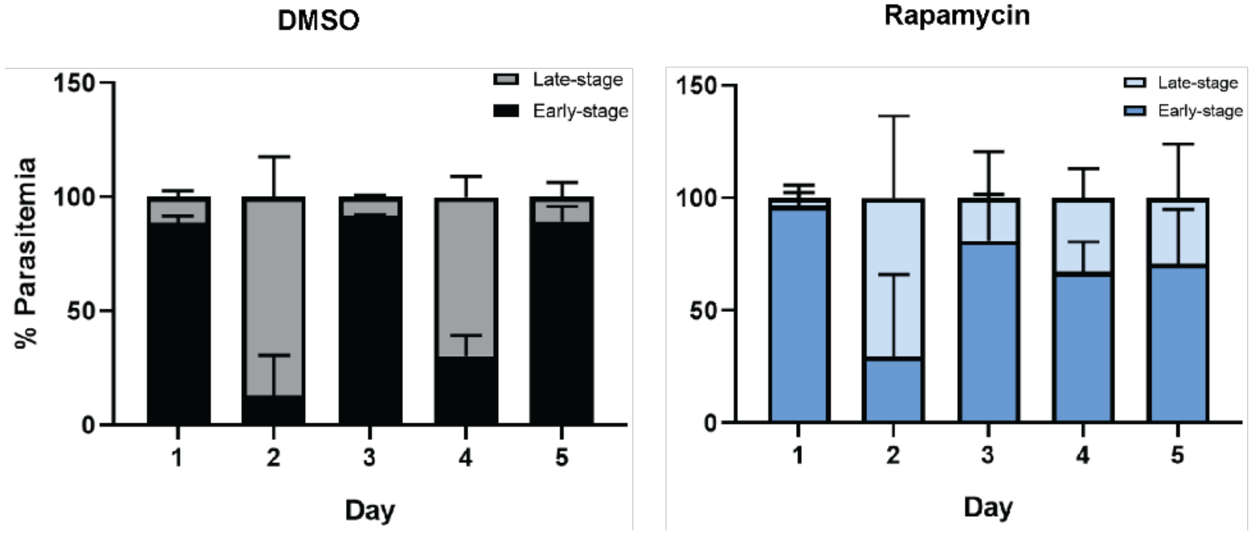


**Figure S2.** Distribution of early- and late-stage parasites in wildtype (DMSO) and K13-mislocalised (Rapamycin) parasites in 2.5 life cycles. Parasitaemia was counted in images of Giemsa-stained uninfected and infected RBCs.

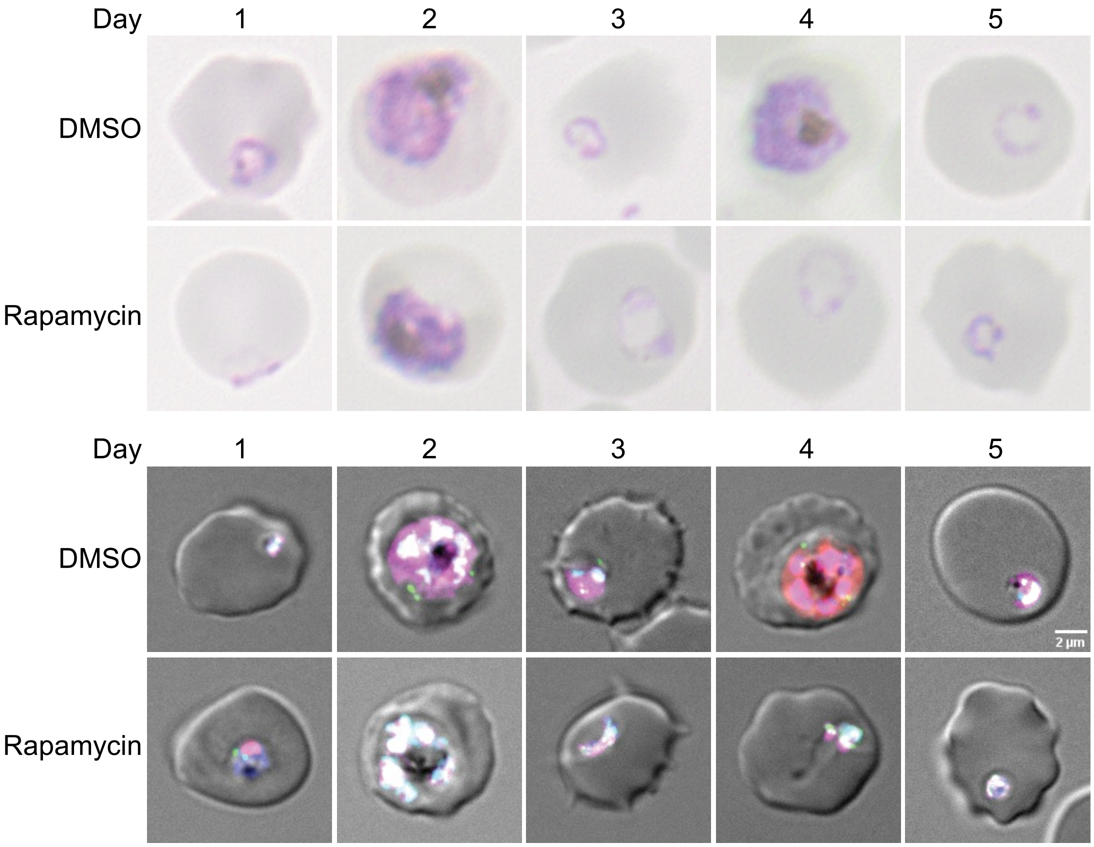


**Figure S3.** Representative images of the Giemsa-stained (top panel) and live-cell images (bottom panel) of K13-100KS-infected RBCs from the mislocalisation assays described in Figure S2. Live-cell images are merged images of DIC (single Z-slice) and fluorescence channels (maximum intensity projections). Cyan (nucleus, Hoechst 33342-stained), magenta (mCherry-NLS), green (GFP-K13).

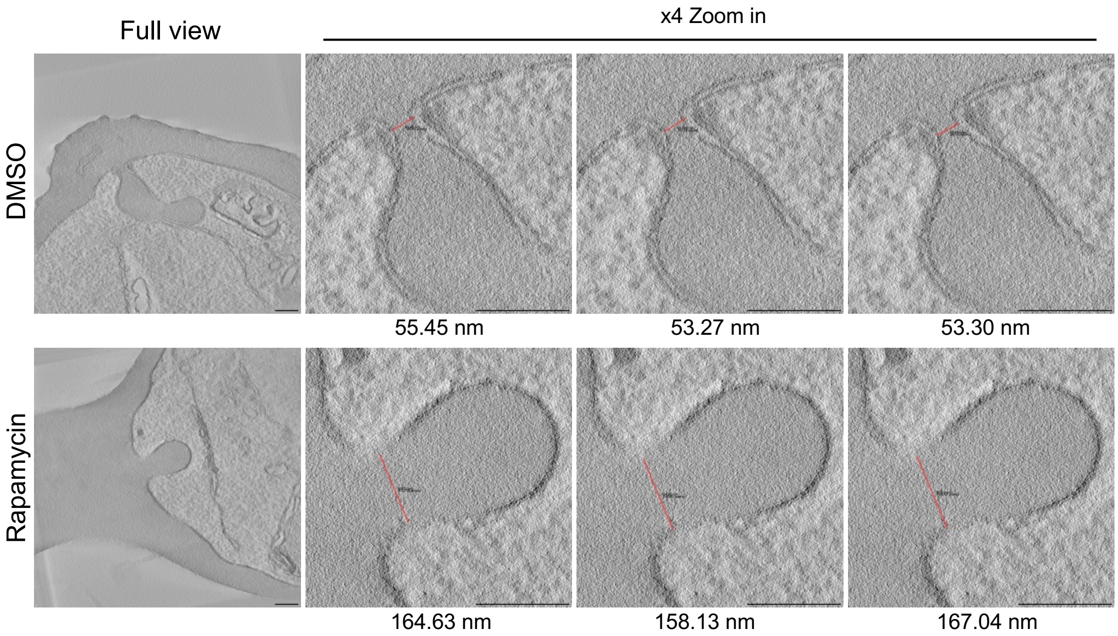


**Figure S4.** Measurements of the neck (tightest or narrowest area in the Z-slice showing the widest opening (approximately neck diameter) to the RBC cytosol) of the invagination of DMSO- and rapamycin-treated K13-100KS-infected RBCs imaged using electron tomography.

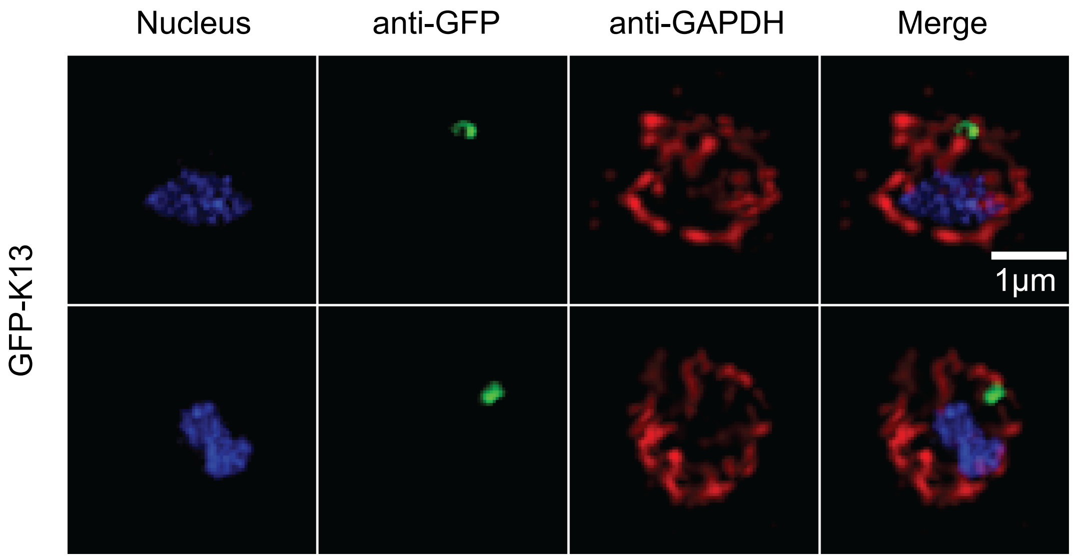


**Figure S5.** Additional images for Figure 1B. IFA of GFP-K13 parasite-infected RBCs imaged using 3D-SIM. Images displayed are maximum intensity projections.

**Table S1.** Parasitemia of the wildtype (DMSO) and K13-mislocalised (Rapamycin) parasites presented in **Figure S2**.

**Raw % parasitemia**

| DMSO | | | | | |
|---|---|---|---|---|---|
| **Early** | | | **Late** | | |
| 10.71 | 5.61 | 6.12 | 1.69 | 0.74 | 0.54 |
| 4.02 | 0.00 | 0.29 | 8.12 | 6.07 | 4.43 |
| 41.92 | 18.95 | 19.85 | 4.32 | 1.59 | 1.80 |
| 11.76 | 16.24 | 13.59 | 45.41 | 33.33 | 22.22 |
| 95.24 | 48.76 | 51.30 | 21.14 | 3.87 | 3.45 |
| **Rapamycin** | | | | | |
| **Early** | | | **Late** | | |
| 11.54 | 6.76 | 5.75 | 0.00 | 0.76 | 0.00 |
| 15.04 | 0.00 | 0.96 | 6.35 | 10.22 | 4.17 |
| 21.00 | 10.65 | 9.35 | 14.51 | 1.95 | 0.00 |
| 26.91 | 11.28 | 10.20 | 24.55 | 3.91 | 3.25 |
| 33.52 | 24.00 | 20.97 | 43.98 | 4.93 | 3.29 |

**Data presented in Figure S2**

| DMSO | | | | | |
|---|---|---|---|---|---|
| **Early** | | | **Late** | | |
| 86.41 | 88.38 | 91.89 | 13.59 | 11.62 | 8.11 |
| 33.10 | 0.00 | 6.12 | 66.90 | 100.00 | 93.88 |
| 90.65 | 92.27 | 91.68 | 9.35 | 7.73 | 8.32 |
| 20.58 | 32.76 | 37.95 | 79.42 | 67.24 | 62.05 |
| 81.84 | 92.64 | 93.70 | 18.16 | 7.36 | 6.30 |
| **Rapamycin** | | | | | |
| **Early** | | | **Late** | | |
| 100.00 | 89.92 | 100.00 | 0.00 | 10.08 | 0.00 |
| 70.30 | 0.00 | 18.68 | 29.70 | 100.00 | 81.32 |
| 59.14 | 84.54 | 100.00 | 40.86 | 15.46 | 0.00 |
| 52.29 | 74.27 | 75.82 | 47.71 | 25.73 | 24.18 |
| 43.25 | 82.97 | 86.43 | 56.75 | 17.03 | 13.57 |

**Table S2.** Raw and processed haem fractionation data for the hemozoin fraction.

| **Early-stage** | | | | | | | | | | | |
|---|---|---|---|---|---|---|---|---|---|---|---|
| **Raw** | | | | | | | | | | | |
| **Sample** | | **DMSO** | | | | | **Rapamycin** | | | | |
| **Biological** | | 1 | 2 | 3 | 4 | 5 | 1 | 2 | 3 | 4 | 5 |
| **Technical** | 1 | 3.95 | 3.51 | 3.43 | - | 4.98 | 4.10 | 3.51 | 3.10 | - | 4.32 |
| | 2 | 4.12 | 3.48 | 3.25 | 4.76 | 4.38 | 4.01 | 3.37 | 3.02 | 4.12 | 4.10 |
| | 3 | - | 3.49 | 3.25 | 4.62 | - | - | 3.36 | 3.13 | 4.05 | - |
| **Average** | | 4.04 | 3.49 | 3.31 | 4.69 | 4.68 | - | - | - | - | - |
| **Normalised to average of DMSO** | | | | | | | | | | | |
| **Sample** | | **DMSO** | | | | | **Rapamycin** | | | | |
| **Biological** | | 1 | 2 | 3 | 4 | 5 | 1 | 2 | 3 | 4 | 5 |
| **Technical** | 1 | 0.98 | 1.01 | 1.04 | - | 1.06 | 1.02 | 1.00 | 0.94 | - | 0.92 |
| | 2 | 1.02 | 1.00 | 0.98 | 1.02 | 0.94 | 0.99 | 0.96 | 0.91 | 0.88 | 0.88 |
| | 3 | - | 1.00 | 0.98 | 0.98 | - | - | 0.96 | 0.95 | 0.86 | - |
| **Average of each biological replicate** | | | | | | | | | | | |
| **Sample** | | **DMSO** | | | | | **Rapamycin** | | | | |
| **Biological** | | 1 | 2 | 3 | 4 | 5 | 1 | 2 | 3 | 4 | 5 |
| | | 1.00 | 1.00 | 1.00 | 1.00 | 1.00 | 1.00 | 0.98 | 0.93 | 0.87 | 0.90 |

| **Late-stage** | | | | | | | | | | | |
|---|---|---|---|---|---|---|---|---|---|---|---|
| **Raw** | | | | | | | | | | | |
| **Sample** | | **DMSO** | | | | | **Rapamycin** | | | | |
| **Biological** | | 1 | 2 | 3 | 4 | 5 | 1 | 2 | 3 | 4 | 5 |
| **Technical** | 1 | 2.33 | 1.73 | 1.16 | 3.86 | 3.78 | 0.83 | 0.65 | 0.39 | 2.18 | 3.28 |
| | 2 | 2.13 | 1.33 | 1.69 | 3.83 | 3.79 | 0.95 | 0.81 | 0.35 | 2.59 | 3.33 |
| | 3 | | 2.00 | 1.12 | 3.87 | 3.75 | 0.97 | 0.66 | 0.43 | 2.66 | 3.12 |
| **Average** | | 2.23 | 1.69 | 1.32 | 3.85 | 3.77 | | | | | |
| **Normalised to average of DMSO** | | | | | | | | | | | |
| **Sample** | | **DMSO** | | | | | **Rapamycin** | | | | |
| **Biological** | | 1 | 2 | 3 | 4 | 5 | 1 | 2 | 3 | 4 | 5 |
| **Technical** | 1 | 1.05 | 1.03 | 0.88 | 1.00 | 1.00 | 0.37 | 0.39 | 0.30 | 0.57 | 0.87 |
| | 2 | 0.95 | 0.79 | 1.28 | 0.99 | 1.00 | 0.43 | 0.48 | 0.26 | 0.67 | 0.88 |
| | 3 | - | 1.19 | 0.84 | 1.00 | 0.99 | 0.43 | 0.39 | 0.32 | 0.69 | 0.83 |
| **Average of each biological replicate** | | | | | | | | | | | |
| **Sample** | | **DMSO** | | | | | **Rapamycin** | | | | |
| **Biological** | | 1 | 2 | 3 | 4 | 5 | 1 | 2 | 3 | 4 | 5 |
| | | 1.00 | 1.00 | 1.00 | 1.00 | 1.00 | 0.41 | 0.42 | 0.29 | 0.64 | 0.86 |

**Table S3.** Raw and processed haem fractionation data for the haem fraction.

| Early-stage | | | | | | | | | | | |
|---|---|---|---|---|---|---|---|---|---|---|---|
| Raw | | | | | | | | | | | |
| Sample | | DMSO | | | | | Rapamycin | | | | |
| Biological | | 1 | 2 | 3 | 4 | 5 | 1 | 2 | 3 | 4 | 5 |
| Technical | 1 | 3.05 | 0.60 | 0.56 | - | 1.58 | 1.23 | 0.45 | 0.47 | 1.03 | - |
| | 2 | 3.01 | 0.53 | 0.58 | 1.51 | 1.47 | 1.27 | 0.50 | 0.48 | - | 0.82 |
| | 3 | - | 0.58 | - | 1.42 | - | - | - | 0.47 | 1.12 | 0.89 |
| Average | | 3.03 | 0.57 | 0.57 | 1.46 | 1.52 | - | - | - | - | - |
| Normalised to average of DMSO | | | | | | | | | | | |
| Sample | | DMSO | | | | | Rapamycin | | | | |
| Biological | | 1 | 2 | 3 | 4 | 5 | 1 | 2 | 3 | 4 | 5 |
| Technical | 1 | 1.01 | 1.05 | 0.99 | - | 1.03 | 0.40 | 0.79 | 0.83 | 0.71 | - |
| | 2 | 0.99 | 0.93 | 1.01 | 1.03 | 0.97 | 0.42 | 0.88 | 0.84 | - | 0.54 |
| | 3 | - | 1.02 | - | 0.97 | - | - | - | 0.82 | 0.76 | 0.58 |
| Average of each biological replicate | | | | | | | | | | | |
| Sample | | DMSO | | | | | Rapamycin | | | | |
| Biological | | 1 | 2 | 3 | 4 | 5 | 1 | 2 | 3 | 4 | 5 |
| | | 1.00 | 1.00 | 1.00 | 1.00 | 1.00 | 0.41 | 0.83 | 0.83 | 0.73 | 0.56 |

| Late-stage | | | | | | | | | | | |
|---|---|---|---|---|---|---|---|---|---|---|---|
| Raw | | | | | | | | | | | |
| Sample | | DMSO | | | | | Rapamycin | | | | |
| Biological | | 1 | 2 | 3 | 4 | 5 | 1 | 2 | 3 | 4 | 5 |
| Technical | 1 | 0.72 | 0.19 | 0.13 | 0.73 | 0.76 | 0.33 | 0.11 | 0.08 | 0.41 | 0.53 |
| | 2 | 0.83 | 0.26 | 0.15 | 0.75 | 0.80 | 0.39 | 0.12 | 0.09 | 0.45 | 0.56 |
| | 3 | 0.80 | - | - | 0.78 | 0.79 | 0.35 | 0.12 | - | 0.47 | 0.55 |
| Average | | 0.78 | 0.22 | 0.14 | 0.75 | 0.78 | - | - | - | - | - |
| Normalised to average of DMSO | | | | | | | | | | | |
| Sample | | DMSO | | | | | Rapamycin | | | | |
| Biological | | 1 | 2 | 3 | 4 | 5 | 1 | 2 | 3 | 4 | 5 |
| Technical | 1 | 0.92 | 0.82 | 0.96 | 0.97 | 0.97 | 0.42 | 0.50 | 0.59 | 0.54 | 0.67 |
| | 2 | 1.06 | 1.18 | 1.04 | 0.99 | 1.02 | 0.50 | 0.51 | 0.65 | 0.60 | 0.71 |
| | 3 | 1.02 | - | - | 1.04 | 1.01 | 0.44 | 0.51 | - | 0.62 | 0.70 |
| Average of each biological replicate | | | | | | | | | | | |
| Sample | | DMSO | | | | | Rapamycin | | | | |
| Biological | | 1 | 2 | 3 | 4 | 5 | 1 | 2 | 3 | 4 | 5 |
| | | 1.00 | 1.00 | 1.00 | 1.00 | 1.00 | 0.45 | 0.51 | 0.62 | 0.59 | 0.69 |
